# Anti-Matrix Protein 1 Monoclonal Antibody Neutralizes Influenza A Virus Subtypes

**DOI:** 10.1101/2025.06.09.658662

**Authors:** Ozlem G. Sahin, Jianhua Le, Rene Devis, Doris J. Bucher

**Affiliations:** Department of Pathology, Microbiology & Immunology, New York Medical College, Valhalla, NY, USA; Junshi Biosciences Co Ltd, Shanghai, China

**Keywords:** Influenza A virus (IAV), Influenza B virus (IBV), H1N1 subtype, H3N2 subtype, Monoclonal antibody, Matrix protein 1 (M1), C-terminal domain (CTD)

## Abstract

**Background:** Research on monoclonal antibodies (mAb) targeting conserved internal proteins of influenza is limited. The matrix protein 1 (M1), the most abundant and conserved internal protein, serves as an endoskeleton bridging cytoplasmic tails of envelope glycoproteins haemagglutinin (HA), neuraminidase (NA) and matrix protein 2 (M2) with viral ribonucleoprotein particles (vRNPs). Clinical studies reveal significant M1 antibody responses post-infection and vaccination, with demonstrated B and T cell recognition. Our study examines 2B-B10-G9, our lab-synthesized mAb targeting conserved linear epitope of M1 at the C-terminal domain (CTD).

**Methods:** Binding of 2B-B10-G9 to the purified influenza A viruses (IAV) and influenza B viruses (IBV) were assessed using SDS-PAGE and Western blotting with Image J analysis. Purified viruses included IAV (H1N1, Pandemic (H1N1) 2009 (H1N1pdm09), and H3N2 subtypes) and IBV which was first isolated in 1940 (B/Lee/40), and B/Victoria lineage. Cytotoxicity of 2B-B10-G9 on the Madin-Darby canine kidney (MDCK) cells was studied by MTT (3-(4, 5-dimethylthiazol-2)-2, 5-diphenyltetrazolium bromide) cell proliferation assay to establish *in vitro* and *in ovo* treatment doses. *In ovo* studies involved injecting 2B-B10-G9 into the allantoic cavity of specific pathogen free (SPF) 10-day-old embryonated chicken eggs (ECE) immediately post-infection with influenza A/Puerto Rico/8/1934 (H1N1) followed by RT-qPCR analysis of the virulence genes HA, NA, and PB1 expressions in the allantoic fluid (AF) 48 hours post-infection.

**Results:** mAb 2B-B10-G9 shows significantly stronger binding to the M1 protein of H1N1 subtype of IAV with respect to H3N2 subtype of IAV (p < 0.001), and it has significantly stronger binding to the M1 protein of H1N1 subtype of IAV with respect to H1N1pdm09 subtype, IBV including B/Lee/40, B/Victoria lineage. (p< 0.0001) mAb 2B-B10-G9 at dose of 40 µg/ml which was determined as an optimal dose according to the MTT assay, significantly reduced plaque-forming units (PFU)/ml of IAV H1N1 and H3N2 subtypes *in vitro* (p< 0.0001). *In ovo* treatment of IAV H1N1 subtype at the same dose 40 µg/ml, significantly suppressed the virulence genes HA, NA, and PB1 expressions compared to the untreated group (p< 0.0001, p< 0.001, p< 0.0001 respectively) and mouse isotype IgG1 mAb treated groups.(p< 0.01)

**Conclusions:** These findings highlight that anti-M1 mAb 2B-B10-G9, targeting conserved linear epitope at the CTD of M1 protein, might be considered as a possible therapeutic option for IAV H1N1 and H3N2 subtypes related to its significant recognition and binding, viral neutralization and suppression of expression of virulence genes including HA, NA and PB1.

## Background

Since the first isolation of the influenza virus in the 1940s, extensive research and public health measures have been implemented to control the spread and impact of influenza^1^. Vaccines and small-molecule antiviral drugs have played critical roles in mitigating outbreaks and treating infected individuals.^1^ Nevertheless, seasonal epidemics and occasional pandemics continue to emerge, posing significant medical and economic burdens to modern societies.^2,3^ One of the major challenges in influenza control is the high mutation rate of circulating strains, driven by antigenic drift and shift in the viral surface glycoproteins hemagglutinin (HA) and neuraminidase (NA).^4,5^ This antigenic variability often undermines the effectiveness of vaccines and therapeutics that target these surface proteins.^5^

To address this limitation, increasing attention has been directed toward more conserved internal viral components as alternative therapeutic targets^6^. Among these, the matrix protein 1 (M1) stands out as the most abundant and conserved structural protein within influenza A (IAV) and B (IBV) viruses. M1, along with matrix protein 2 (M2), is encoded by RNA segment 7 and produced via alternative splicing^7^. While IAV and IBV M1 proteins share only ∼30% sequence identity, both are essential for viral assembly and budding^7^. M1 of IAV comprises 252 amino acids, while IBV M1 contains 248 amino acids ^7, 8^. The C-terminal domain (CTD) of M1 plays a particularly critical role by interacting with viral ribonucleoproteins (vRNPs), regulating their intracellular trafficking, and facilitating viral egress through interactions with host membranes and induction of membrane tubulation ^9,10^.

Resistance to antiviral drugs, especially in IAV, led to the discontinuation of M2 proton channel inhibitors and the adoption of NA inhibitors (oseltamivir, zanamivir, peramivir) and RdRP inhibitors (baloxavir marboxil).^11,12^ However, antiviral resistance particularly in immunocompromised patients and antigenic drifts of the viral surface proteins targeted by monoclonal antibodies (mAbs) remain a major concern in prophylactic and therapeutic approaches for influenza ^13–17^.

In this study, we report the characterization of a hybridoma cell line, 2B-B10-G9, which secretes a monoclonal antibody (mAb) targeting a conserved linear epitope within the CTD of M1. This antibody demonstrated strong binding affinity and in vitro neutralization activity against both H1N1 and H3N2 IAV subtypes. Additionally, it significantly suppressed the expression of key virulence genes, including HA, NA, and PB1, in H1N1-infected embryonated eggs (in ovo). These findings highlight the therapeutic potential of targeting the conserved M1 protein, and specifically its CTD, offering a promising alternative or complement to existing influenza treatments that primarily focus on the mutable surface glycoproteins.

## METHODS

### Cells and Viruses

MDCK II epithelial cells (Madin-Darby canine kidney, ATCC, Manassas, VA, USA) were cultured in T75 flasks with Minimum Essential Medium (1X MEM) (Gibco) supplemented with 20% heat-inactivated FBS, 10 U/ml penicillin, 10 µg/ml streptomycin, 5% sodium bicarbonate, 1.0 M HEPES, 0.2 M L-glutamine (all from Sigma-Aldrich), and autoclaved deionized H_2_O. Cells were maintained at 37°C in 5% CO_2_. For subculture, cells at ∼90% confluence were dissociated using 0.05% Trypsin-EDTA (Sigma-Aldrich) and seeded into culture dishes at 500,000 cells/dish in 8 ml 1X MEM, growing until ∼90% confluence. The following IAV and IBV were propagated in 10-day-old embryonated chicken eggs and purified by ultracentrifugation per established protocols.^18, 19^ (Table 1)

**Table 1.** Influenza virus subtypes/lineages and strains used in our studies (*in vitro* and *in ovo*)

| Virus | Subtype/Lineage | Strain | Study |
| --- | --- | --- | --- |
| IAV | H1N1 | A/Puerto Rico/8/1934 | *In vitro, In ovo* |
| IAV | H1N1pdm09 | A/California/07/2009 | *In vitro* |
| IAV | H1N1pdm09 | A/Victoria/2570/2019 | *In vitro* |
| IAV | H3N2 | A/Uruguay/716/2007 | *In vitro* |
| IBV | Prototype | B/Lee/40 | *In vitro* |
| IBV | Victoria | B/Austria/1359417/2021 | *In vitro* |

IAV: Influenza A virus. IBV: Influenza B virus.

Acceptable titers for *in vitro* and *in ovo* infections were determined using MDCK plaque assays and 50% egg infective dose (EID_50_)/ml, as described. ^20, 21^

### Generation of hybridoma cell line for the anti-matrix protein 1 mAb 2B-B10-G9

Hybridoma cell lines used in this study were generated by fusing pooled splenocytes from 12-week-old female BALB/c mice, following the method described by Bucher et al.^22^ **Immunization Protocol:** Mice were immunized intraperitoneally on days 0, 14, and 28 with 25 μg of purified M1 protein from A/Puerto Rico/8/1934 (H1N1) in 200 µL of PBS, mixed with 100 µL of complete Freund’s adjuvant (CFA). On day 40, blood samples were collected, and sera were tested for M1 protein response via ELISA. On day 45, mice received a final intravenous boost of 25 μg purified M1 protein in 200 µL of PBS with 100 µL of CFA via the tail vein.

### Hybridoma Generation

On day 48, mice were euthanized, and their spleens were removed and washed with PBS. Spleen cell suspensions were prepared by pressing the spleens through a cell strainer into 20 mL of cell culture medium. Splenocytes were fused with X63-Ag 8.6539 mouse myeloma cells at a 1:3 ratio using pre-warmed polyethylene glycol, following the Kohler and Milstein protocol.^23^ Fused cells were cultured in 2 mL of hypoxanthine-aminopterin-thymidine (HAT) medium in 24-well plates for 10–14 days. Supernatants from wells producing monoclonal antibodies (mAbs) against M1 protein (confirmed via ELISA) were selected for antibody purification.

### Antibody purification by protein G column chromatography

Antibody purification was performed using Protein G column chromatography on cell culture supernatants from the 2B-B10-G9 hybridoma cell line, following the method previously described.^24^

### The Lowry method for protein quantification

Ten dilutions of 200 μL protein standard and unknown protein samples (purified anti-M1 mAb 2B-B10-G9, purified IAV, and IBV) were prepared in test tubes. Protein quantification was performed using Lowry reagent (Sigma-Aldrich, St. Louis, MO, USA) and Folin-Ciocalteu reagent (Sigma-Aldrich, St. Louis, MO, USA), following the method previously described.^25^

### SDS-PAGE and staining with Coomassie brilliant blue R250 of the anti-matrix protein 1 mAb 2B-B10-G9 and mouse isotype IgG1 mAb

A total of 1.00 μg of mAb 2B-B10-G9 and mouse isotype IgG1 mAb were mixed with 5 μL NuPAGE® lithium dodecyl sulfate (LDS) sample buffer (4x) (Invitrogen, Carlsbad, CA, USA), 2 μL NuPAGE® sample reducing agent (10x) (Invitrogen, Carlsbad, CA, USA), and deionized H2O to bring the final volume to 20 μL. The samples were heated at 70 °C for 10 minutes in a water bath. The prepared samples were loaded onto NuPAGE® Novex 4-12% Bis-Tris precast gel (Invitrogen, Carlsbad, CA, USA) and separated using the Invitrogen XCell SureLock mini-cell electrophoresis system at 200V constant for 30 minutes in NuPAGE® MES SDS running buffer (Invitrogen, Carlsbad, CA, USA). After electrophoresis, the gel was treated with a solution of 10% acetic acid, 50% methanol, and 40% H2O for 3 hours with shaking, changing the solvent every hour to effectively remove SDS from the gel. The gel was then stained with 0.1% Coomassie Blue R250 in 10% acetic acid, 50% methanol, and 40% H2O for 1 hour with shaking. Following staining, the gel was de-stained by shaking in 10% acetic acid, 50% methanol, 40% H2O for 2 hours, with the solvent being changed every hour.^26^

### SDS-PAGE and Western blotting of the anti-matrix protein 1 mAb 2B-B10-G9 with the purified influenza A and influenza B viruses

10 μg each of purified IAV and IBV and 5 μL of deionized H2O were mixed with 5 μL NuPAGE® LDS Sample Buffer (4x) (Invitrogen, Carlsbad, CA, USA) to bring the final volume to 20 μL. The samples were heated at 70 °C for 10 minutes in a water bath, then loaded onto NuPAGE® Novex 4-12% Bis-Tris precast gel (Invitrogen, Carlsbad, CA, USA). Samples were separated using the Invitrogen XCell SureLock mini-cell electrophoresis system at 200V constant for 30 minutes in NuPAGE® MES SDS running buffer (Invitrogen, Carlsbad, CA, USA). Upon completion of electrophoresis, the gels were transferred to polyvinylidene difluoride (PVDF) membrane for Western blot analysis as previously described.^27^

### Quantification of the protein bands with the Image J algorithm

The binding of mAb 2B-B10-G9 to the M1 protein of A/Puerto Rico/8/1934 (H1N1), B/Lee/40, B/Austria/1359417/2021 (B/Victoria), A/California/07/2009 (H1N1pdm09), A/Victoria/2570/2019 (H1N1pdm09) and A/Uruguay/716/2007 (H3N2) were quantified from Western blots using the ImageJ algorithm as follows:

1. Protein intensities (PI) were measured from each rectangular band on the blot. A dot (+) was placed at the center of each rectangular band.
2. The inverted PI was calculated by subtracting the PI from the highest pixel intensity value, 255.
3. Background intensity (BI) was measured adjacent to each protein band. The inverted BI was calculated by subtracting the BI from 255.
4. The loading control allantoic fluid (AF) band and dot (+) intensities (AFI) were measured, and the inverted AFI was calculated by subtracting AFI from 255.
5. The net PI was calculated by subtracting the inverted BI from the inverted PI.

The mean values of the net PI/net AFI ratios of band and dot intensities were used for the quantification of the protein bands.^28^

### MTT Assay: quantification of cell viability

MTT powder was dissolved in Dulbecco’s Phosphate Buffered Saline, pH=7.4 (DPBS) to 5 mg/ml. MTT solution was filter-sterilized through a 0.2 µM filter into a sterile, light protected container and stored at 4°C for frequent use or at -20°C for long term storage. MDCK cells were seeded in 96-well plate (∼10,000 cells/well) and mAb 2B-B10-G9 at concentrations of 0.5, 1.0, 5.0, 10.0, 40.0, 90.0 and 200 ug/ml containing a final volume of 100 µl/well were added in three technical replicates. Staurosporine, an apoptotic agent at 1µM concentration, was used as a positive control. DMSO (0.5 %) was used as a vehicle (negative) control. After 24 hours of incubation at 37°C, 5% CO_2_, 10 µl MTT solution (5mg/mL) was added to each well, and incubated for 4 hours at 37°C, 5% CO_2_ by protection from light. After 4 hours of incubation, 150 µl solubilization solution (100% DMSO) was added to each well to dissolve formazan crystals. The 96-well plate was wrapped in foil and shaked on an orbital shaker for 15 minutes to dissolve the formazan crystals. The absorbance/OD at 590 nm was recorded within 1 hour. (Results demonstrated in the supplement file)

### Plaque assay: quantification of plaque forming unit/ml (PFU/ml)

MDCK cells were seeded in cell culture dishes (Corning, Hickory, NC, USA) with 1X MEM. Upon reaching approximately 90% confluency, the media was removed, and the cells were washed with 2 mL of cell wash/diluent solution containing 1X PBS, 0.2% bovine serum albumin (BSA) (Sigma-Aldrich, St. Louis, MO, USA), and 1% calcium-magnesium (bivalent cations). Serial dilutions of PR8, H1N1 and A/Uruguay/716/2007 (H3N2) subtypes were prepared in three technical replicates at 10^−2^, 10^−3^, 10^−4^ dilutions using the cell wash/diluent solution. 400 µL of each virus suspension was added to each cell culture dish. The dishes were incubated for 60 minutes at 37°C with intermittent rocking (at 30 minutes), in the presence of 5% CO_2_. Following incubation, the virus inoculum was removed. An agarose overlay was prepared consisting of 2X MEM, 0.4% BSA, half the volume of autoclaved deionized H2O, 2% agarose (Seakem LE, Lonza Bioscience, Morrisville, NC, USA), and 2.5 µL of TPCK-treated trypsin (Fisher Scientific, Pittsburgh, PA, USA) (2 mg/mL stock, final concentration: 2 µg/mL). 8 mL of this overlay was added to each cell culture dish, and the dishes were incubated for 72 hours post-infection. After 72 hours, the agar plugs were removed, and the virus-infected MDCK monolayers were stained with 0.1% crystal violet in 20% ethanol. Plaques were counted. The plaque-forming units per milliliter (PFU/mL) were calculated using the formula: PFU/mL= N×V/D N = Average number of plaques in three technical replicate dishes D = Dilution factor of the virus V = Volume of the virus inoculated (in mL) For the plaque reduction neutralization (PRNT) assay, the dilution factor of 10^−4^ for PR8, H1N1 and A/Uruguay/716/2007 (H3N2) forming 60-70 countable plaques was chosen as demonstrated in Figure 1 for A/Uruguay/716/2007 (H3N2).

**Figure 1.**
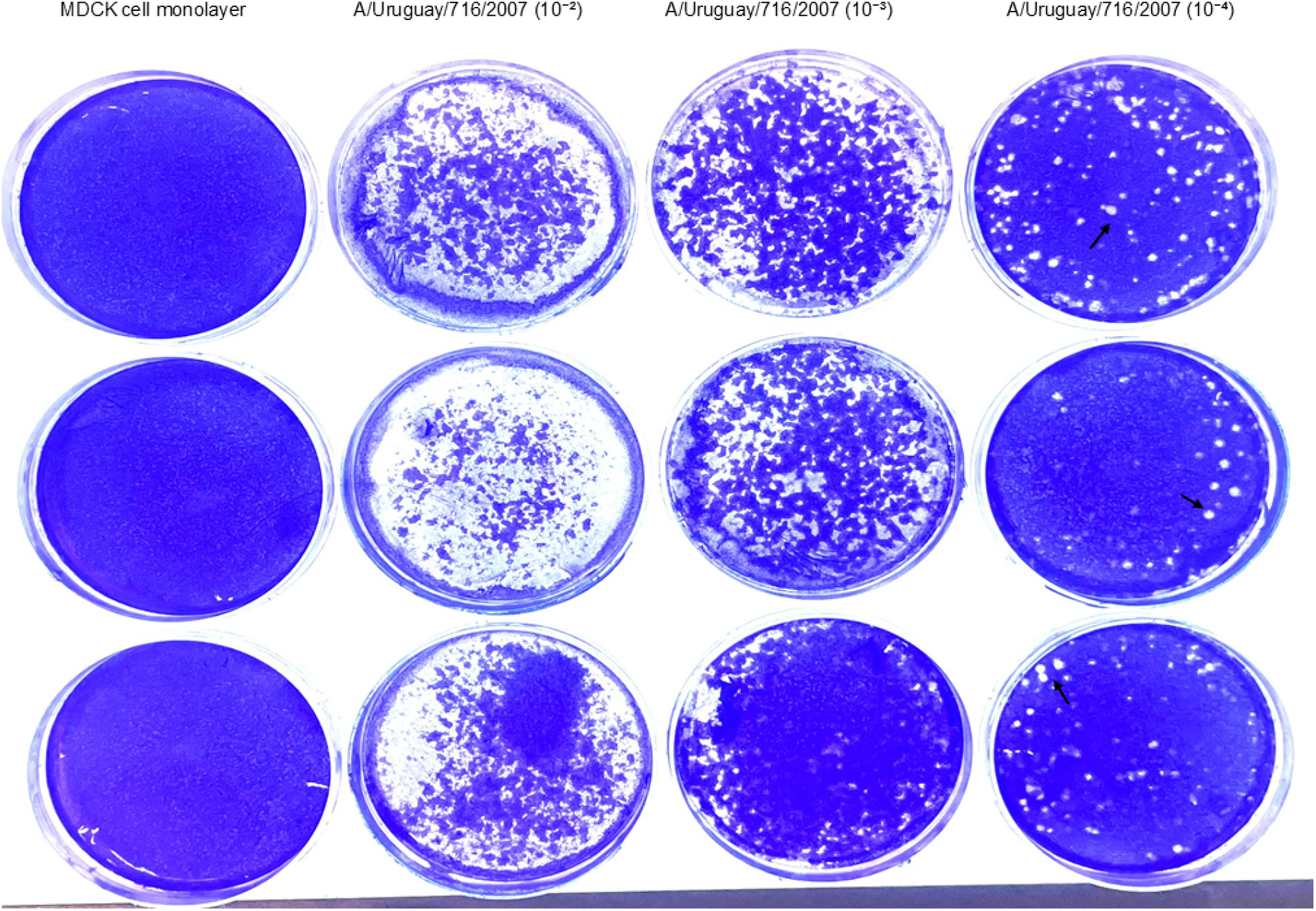
Plaque assay of A/Uruguay/716/2007 (H3N2) grown in the MDCK cell for determination of dilution factor for neutralization test. First column of dishes is control cells with no virus in three technical replicates. Columns 2-4 are serial dilutions (10^−2^, 10^−3^ and 10^−4^) of A/Uruguay/716/2007 (H3N2) Plaques are indicated by black arrows. This experiment set was repeated for three biological replicates of different MDCK cell lines leading to identical conclusions.

### Plaque reduction neutralization test (PRNT)

PR8, H1N1 and A/Uruguay/716/2007 (H3N2) from the viral stocks were diluted to 1.6 × 10^6^ PFU/mL in the cell wash/diluent solution. 400 µL of this viral suspension, approximately 65 PFU of virus, was added to each designated cell culture dish of MDCK cells grown to ∼90% confluency. Prior to virus inoculation, the cells were washed with 4 mL of cell wash/diluent solution. Cell culture dishes were prepared in three technical replicates for the treatment doses of mAb 2B-B10-G9. Pre-treatment dishes received the virus inoculum without treatment. Post-treatment dishes received virus inoculum and treated with mAb 2B-B10-G9.The cell culture dishes were rocked briefly and incubated for 60 minutes at 37°C with 5% CO_2_, with intermittent rocking at 30 minutes. After the incubation, the viral suspension was aspirated, and 100 µL of mAb 2B-B10-G9 treatment (at 40.0 µg/mL in 8 mL 2xMEM) was added as three technical replicates. The dishes were then covered with the complete agarose overlay and incubated inverted for 72 hours at 37°C with 5% CO_2_. After the incubation period, the agar plugs were removed, and the MDCK cell monolayers were stained with 0.1% crystal violet in 20% ethanol. The plaques from the three technical replicates were counted. The average number of plaques and PFU/ml were calculated as previously described.^29^

### *In ovo* inoculation of A/Puerto Rico/8/1934 (PR8, H1N1)

100 µl of PR8, H1N1 at 10^−7^ dilution (EID50) was injected into the allantoic cavity of 10-day-old specific pathogen-free embryonated chicken eggs. Immediately after viral infection, 100 µL of mAb 2B-B10-G9 was injected into the allantoic cavity having 8 ml of allantoic fluid (AF) with the final mAb concentration of 40.0 µg/mL.Untreated eggs received the virus injection only, without the mAb 2B-B10-G9 treatment. Uninfected eggs received the mAb 2B-B10-G9 injection without the virus injection. Isotype (IgG1) treated eggs received the virus injection and treated with the mouse isotype IgG1 mAb at 40.0 µg/mL. All eggs were incubated at 37°C for 48 hours after virus inoculation and immediate mAb treatments. After 48 hours of incubation, AF was harvested from each egg for viral RNA extraction. The AF from the uninfected and untreated eggs were also harvested for RNA extraction of the housekeeping gene beta-actin (B-ACT) as an internal control.

### RNA Extraction of A/Puerto Rico/8/1934 (PR8, H1N1) and the housekeeping gene beta actin (B-ACT) with mini spin procedure

RNA extraction for virulence genes the HA, NA, PB1 of PR8, H1N1 and the housekeeping gene B-ACT from the AF were performed with the mini spin procedure according to the manufacturer’s protocol.^30^

### Measurement of the RNA concentration of samples extracted from the allantoic fluid (AF)

NanoDrop ND-1000 spectrophotometer (Thermo Fisher Scientific, Wilmington, DE USA) was used to measure the RNA concentration of samples isolated from AF using 1.0 µl total RNA. Quantification of the RNA was based on measuring ultraviolet absorbance at 260 nm and 280 nm. RNA purity was judged by the 260 nm / 280 nm ratio, and a low ratio was considered as contamination by proteins.^31^

### RT-qPCR for viral virulence genes HA, NA, PB1 and housekeeping B-ACT gene quantification

RNA Samples from the uninfected, untreated, isotype (IgG1) treated, mAb 2B-B10-G9 treated, and egg internal control groups were analyzed using CFX96 Bio-Rad RT-qPCR system (Bio-Rad, Hercules, CA, USA). The experiment was designed as three biological replicates from three different eggs for each gene analysis, and repeated three times on different days. Amplification of the HA, NA, PB1, and B-ACT genes was carried out using the Bio-Rad manufacturer’s protocol (2010). Forward and reverse primers for the HA, NA, PB1 genes of PR8, H1N1 and FAM probe were mixed with 10 mM Tris buffer (TE buffer).^32^ Forward and reverse primers for B-ACT gene and FAM probe were used for normalizing mRNA expression in chicken tissues.^33^ For each gene, a master mix was prepared using the respective probe mixture, nuclease-free water (IDT, Newark, NJ, USA), and RT-qPCR ready mix (Bio-Rad, Hercules, CA, USA). The RT-qPCR ready mix contained murine leukemia virus (MMLV) reverse transcriptase, thermus aquaticus (Taq) DNA polymerase, dNTPs, and MgCl2 for cDNA synthesis. 18 µL of the master mix and 2 µL of RNA sample were added to 96-well PCR plates (USA Scientific, Ocala, FL, USA). The plates were sealed and spun to remove air bubbles using a plate spinner (Corning, Hickory, NC, USA). The RT-qPCR program was set for 40 cycles with denaturation at 95°C for 10 seconds, annealing and polymerization at 62°C for 30 seconds. Normalization was done using quantification cycle (Cq) values, where Cq is the number of cycles required for the FAM fluorescence to reach a threshold.^34^ Δ Cq values were calculated by subtracting the Cq (B-ACT) from the Cq (HA, NA, PB1) for each sample. Δ Δ Cq was calculated by subtracting the average Δ Cq of the uninfected groups from the Δ Cq values of the other groups. Gene expression was determined using the 2^- Δ Δ Cq^ .^35^ Log2 fold change calculation: log2(fold change)=log2(gene expression in condition A)-log2(gene expression in condition B). ^36^

### Statistical analysis

All statistical analyses were performed using Prism software (GraphPad version 10). The number of technical and biological repeats for experiments and specific tests for statistical significance used are indicated in the corresponding figure legends.

## RESULTS

### Anti-matrix protein 1 mAb 2B-B10-G9 is IgG1 isotype

mAb 2B-B10-G9 antibody was generated using hybridoma technology and IgG1 isotype with a total protein concentration of 3.20 mg/ml. SDS-PAGE under reducing conditions stained with Coomassie brilliant blue R250 showed whole mAb 2B-B10-G9 at 150 kDa, heavy chain at 50 kDa, and light chain at 25 kDa regions (Figure 2 lanes 1-4). The mouse monoclonal IgG1 isotype heavy chain was detected at 50 kDa and light chain 25 kDa regions (Figure 2 lane 5).

**Figure 2.**
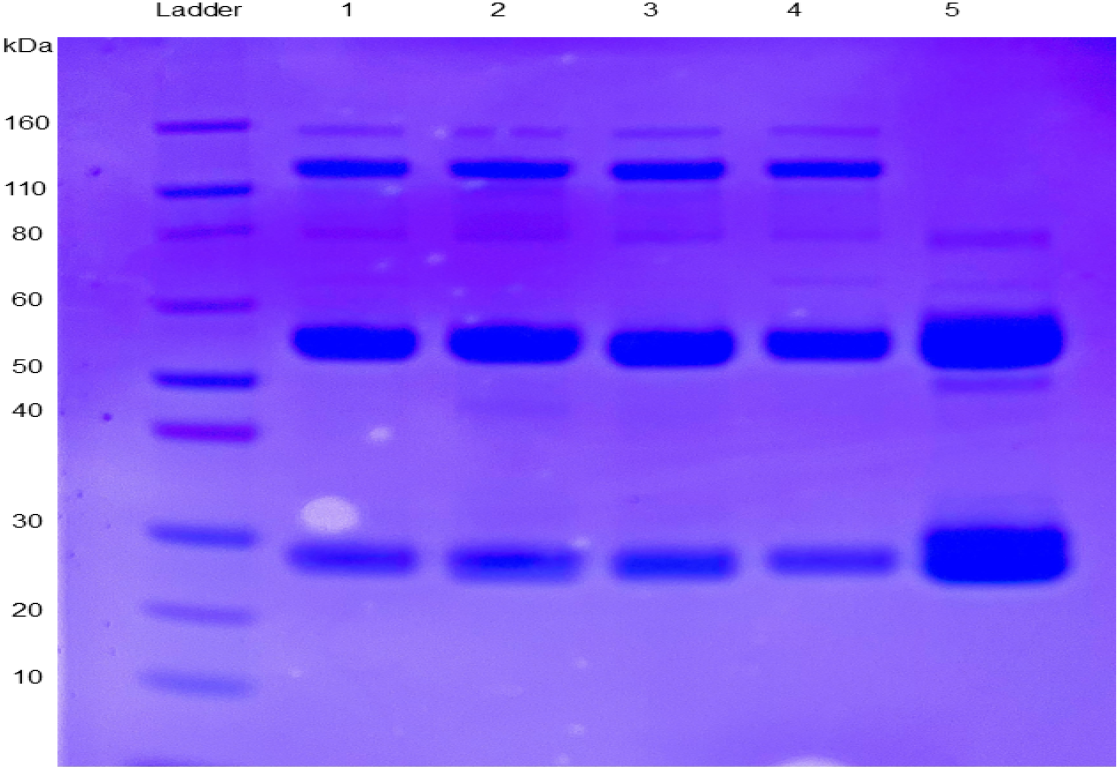
Demonstration that anti-matrix protein 1 mAb 2B-B10-G9 is IgG1 isotype. The figure shows SDS-PAGE under reducing condition and Coomassie brilliant blue staining of mAb 2B-B10-G9 **(Lane 1-4)** and the mouse IgG1 isotype mAb **(Lane 5)**. IgG1 (whole molecule) at **150 kDa** region **(Lane 1-4)**, heavy chain at **50 kDa** region **(Lane 1-5)** and light chain at **25 kDa** region **(Lane 1-5)** were depicted.

### mAb 2B-B10-G9 shows heterosubtypic binding to the matrix protein 1 of influenza A viruses

Western blot data were quantified using the Image J algorithm with the mean of net PI/net AFI ratios calculated from dot and band intensities (see the Materials and Methods section). The mean of net PI/net AFI ratios were highest for A/Puerto Rico/8/1934 (H1N1) followed by A/Uruguay/716/2007 (H3N2), A/Victoria/2570/2019 (H1N1pdm09) and A/California/07/2009 (H1N1pdm09) subtypes (Table 3). The lowest means of net PI/net AFI ratios were observed for B/Lee/40 and B/Austria/1359417/2021 (Table 3). Anti-M1 mAb 2B-B10-G9 showed heterosubtypic binding to the H1N1, H3N2 and H1N1pdm09 subtypes of IAV in the 30 kDa region (Figure 3A). The binding of mAb 2B-B10-G9 to IBV was negligible (Figure 3A). mAb 2B-B10-G9 showed significantly higher binding to the M1 protein for PR8, H1N1 followed by A/Uruguay/716/2007 (H3N2), A/Victoria/2570/2019 (H1N1pdm09) and A/California/07/2009 (H1N1pdm09) subtypes with respect to IBV B/Lee/40 and B/Austria/1359417/2021 (p< 0.0001) (Figure 3B).

**Table 2.**
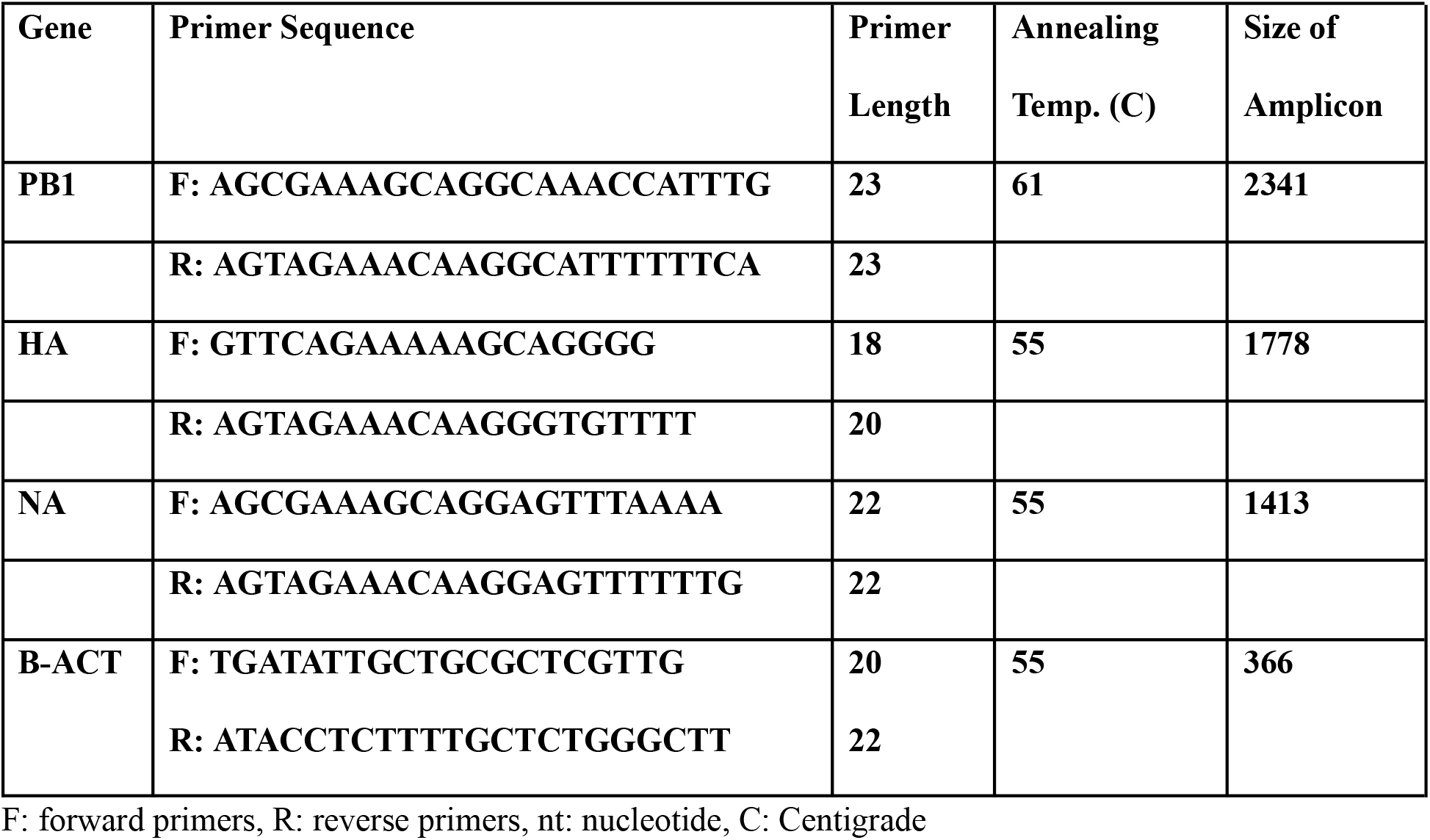
Oligonucleotide Primers used in RT-PCR.

**Table 3:**
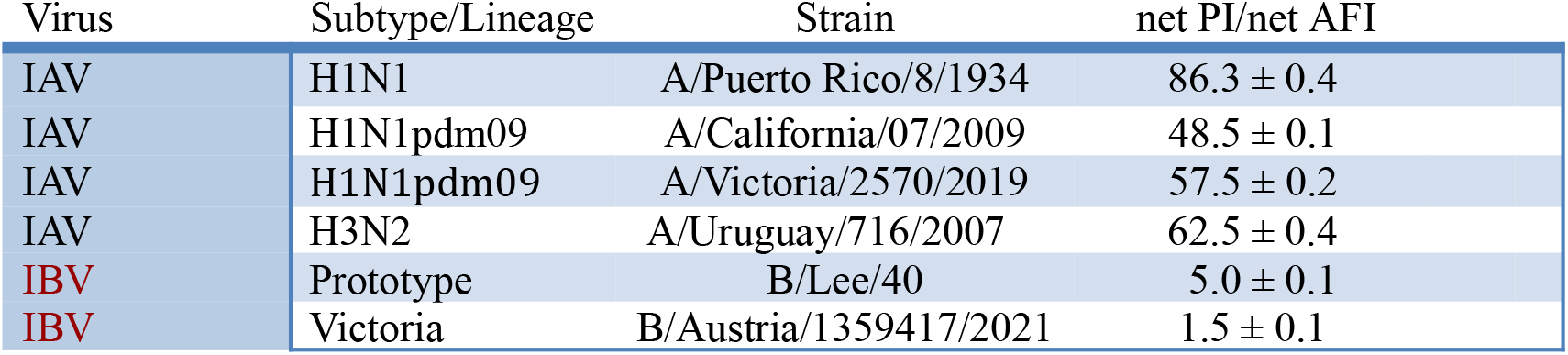
Binding of mAb 2B-B10-G9 to the matrix protein 1 of influenza A and influenza B viruses. Binding is indicated by the ratio of net protein intensity (PI) to the net allantoic fluid intensity. (AFI) High ratios indicate stronger binding. There was strong binding to IAV subtypes in contrast to the weak binding to IBV.

**Figure 3.**
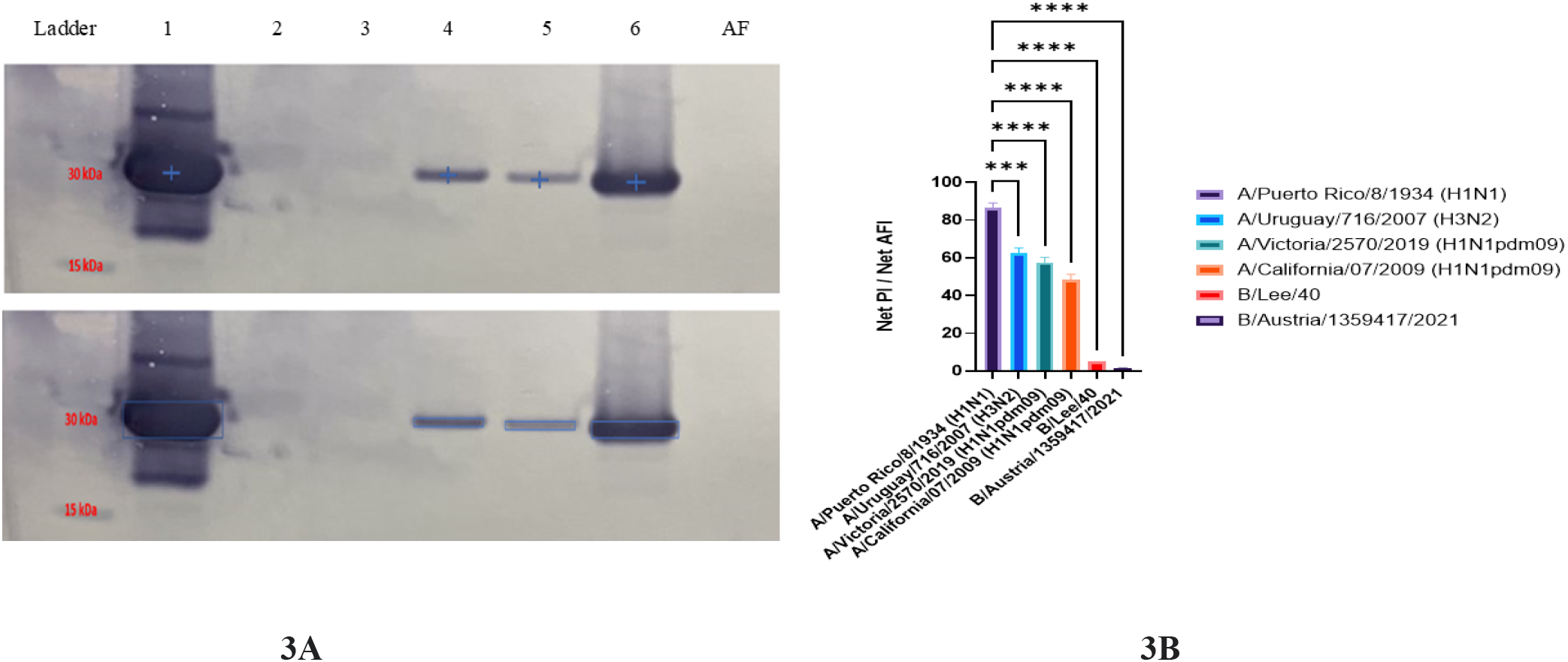
**A. Anti-matrix protein 1 mAb 2B-B10-G9 shows heterosubtypic binding to the H1N1, H3N2 and H1N1pdm09 subtypes of influenza A viruses at 30 kDa region**. Western blots for purified IAV and IBV between lane 1-6 and lane 7 allantoic fluid (AF) as internal control separated by SDS-PAGE under non-reducing conditions and probed with mAb 2B-B10-G9. Lane1: A/Puerto/Rico/8/1934 (PR8, H1N1) Lane 2: B/Lee/40, Lane 3: B/Austria/1359417/2021, Lane 4: A/Victoria/2570/2019 (H1N1 pdm09), Lane 5: A/California/07/2009 (H1N1 pdm09) Lane 6: A/Uruguay/716/2007 (H3N2), Lane 7: AF. **Figure 3B. Anti-matrix protein 1 mAb 2B-B10-G9 shows significant binding to IAV H1N1 subtype, H3N2, and H1N1pdm09 subtypes while binding to IBV was negligible**. 2B-B10-G9 binding to the M1 protein was quantitated with the Image J software algorithm. Data of two technical replicates are represented as mean ± SD. One-Way ANOVA Multiple Comparisons was used to compare the means. ***p < 0.001 ****p< 0.0001.

### Anti-matrix protein 1 mAb 2B-B10-G9 neutralizes H1N1 and H3N2 subtypes of Influenza A virus

Plaque reduction neutralization test (PRNT) was used to assess the efficacy of mAb 2B-B10-G9 at 40 µg/ml in neutralizing H1N1 and H3N2 subtypes of IAV. For A/Puerto Rico/8/1934 (PR8, H1N1) average plaque counts were reduced from 65.0 to 3.6 (Figure 4A). For A/Uruguay/716/2007 (H3N2), average plaque counts were reduced from 56.7 to 6.0 (Figure 4A). Reduction of PFU × 10^5^/ml for A/Puerto Rico/8/1934 (PR8, H1N1) was from 16.8 to 1.6 after mAb 2B-B10-G9 treatment (Figure 4B). Reduction of PFU × 10^5^/ml for A/Uruguay/716/2007 (H3N2) was from 14.2 to 1.5 after mAb 2B-B10-G9 treatment (Figure 4B). mAb 2B-B10-G9 treatment at 40 µg/ml demonstrated significantly potent (90%) reduction in average plaque counts and PFU x10^5^ /ml for IAV H1N1 and H3N2 subtypes (p< 0.0001).

**Figure 4.**
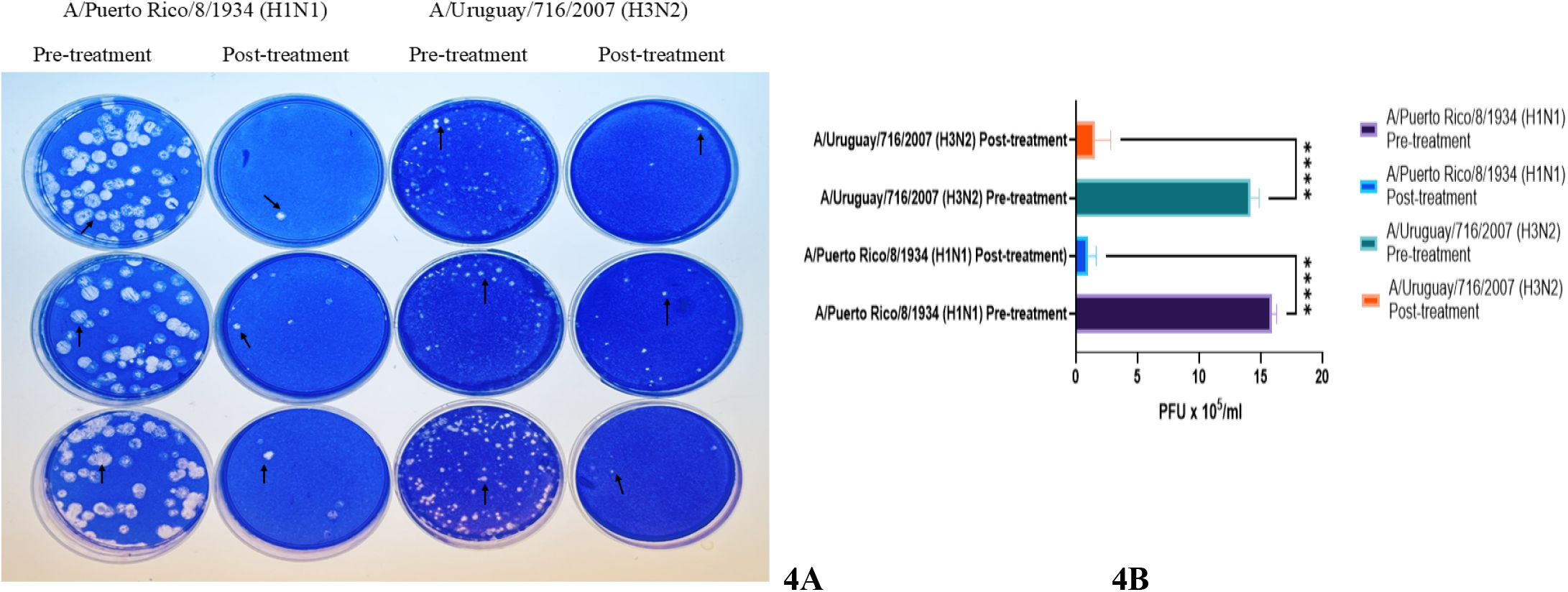
**A. *In vitro* neutralization of A/Puerto Rico/8/1934 (PR8, H1N1) and A/Uruguay/716/2007 (H3N2) by anti-matrix protein 1 mAb 2B-B10-G9 in plaque reduction neutralization test.** *In vitro* neutralizing efficacy of mAb 2B-B10-G9 at 40 µg/ml is represented as reduction in the number of plaques in three technical replicates. Black arrows indicate the plaques. **Figure 4B. *In vitro* neutralizing efficacy of mAb 2B-B10-G9 at 40 µg/ml is significant for reduction in PFU x10**^**5**^ **/ml**. Pre-treatment and post-treatment PFU x10^5^ /ml of A/Puerto Rico/8/1934 (PR8, H1N1) and A/Uruguay/716/2007 (H3N2) from three technical replicates is represented in the bar graph. Experiment was repeated three times with the same results and the data are represented as mean ± SD. One-Way ANOVA Multiple Comparisons was used to compare the means. ****p< 0.0001.

### Anti-matrix protein 1 mAb 2B-B10-G9 suppresses virulence gene expressions of A/Puerto Rico/8/1934 (PR8, H1N1) *in ovo*

mAb 2B-B10-G9 treatment resulted in reduction of virulence gene expressions for the HA, NA, and PB1 genes of PR8, H1N1. The mean ± SD of (2^- Δ Δ Cq) of gene expressions and log2-fold-change (FC) values for the HA, NA and PB1 genes of the untreated, mAb 2B-B10-G9 treated, isotype (IgG1) mAb treated and uninfected groups were summarized in Table 4. The mean values of (2^- Δ Δ Cq) for the HA, NA and PB1 genes in the PR8, H1N1 infected groups were 60924, 284464 and 96374 respectively. (Table 4) Treatment with mAb 2B-B10-G9 decreased the mean values of (2^- Δ Δ Cq) for the HA, NA and PB1 genes to 1.9, 3.2 and 2.9 with log2FC values of -14.9, -16.4 and -15.0 respectively. (Table 4) Treatment with isotype IgG1 mAb decreased the mean values of (2^- Δ Δ Cq) for the HA, NA and PB1 genes to 179, 251 and 307 with log2FC values of -8.4, -10.1 and -8.2 respectively. (Table 4) There was a statistically significant reduction of the HA, NA and PB1 gene expressions of PR8, H1N1 in the mAb 2B-B10-G9 treatment group with respect to the untreated group. (p< 0.0001, p< 0.001 and p< 0.0001 respectively) (Figure 5) There was also statistically significant reduction of the HA, NA and PB1 gene expressions of PR8, H1N1 in the isotype IgG1 mAb treatment group with respect to the untreated group. (p< 0.01, p< 0.05 and p< 0.01 respectively) (Figure 5) However, mAb 2B-B10-G9 treatment group showed statistically significant reduction of the HA, NA and PB1 gene expressions of PR8, H1N1 with respect to the isotype IgG1 mAb treatment group. (p< 0.01) (Figure 5)

**Table 4.** Demonstration that treatment with anti-matrix protein 1 mAb 2B-B10-G9 suppresses virulence gene expressions of A/Puerto/Rico/8/1934. (PR8, H1N1) The table shows the mean ± SD (2^ Δ Δ Cq) and log2foldchange (FC) values for the hemagglutinin (HA), neuraminidase (NA) and polymerase basic 1 (PB1) genes of PR8, H1N1 in the untreated, mAb 2B-B10-G9 treated, isotype IgG1 mAb treated and uninfected groups.

| Groups | HA ($2^{-\Delta\Delta Cq}$) log2FC | NA ($2^{-\Delta\Delta Cq}$) log2FC | PB1 ($2^{-\Delta\Delta Cq}$) log2FC |
| --- | --- | --- | --- |
| Untreated | 60924 ± 90.3 | 284464 ± 43.5 | 96374 ± 14.3 |
| 2B-B10-G9 treatment | 1.9 ± 1.3 -14.9 | 3.2 ± 2.2 -16.4 | 2.9 ± 2.2 -15.0 |
| Isotype (IgG1) treatment | 179 ± 2.9 -8.4 | 251 ± 3.3 -10.1 | 307 ± 5.4 -8.2 |
| Uninfected | 1.1 ± 0.4 -15.7 | 1.0 ± 0.3 -18.1 | 1.1 ± 0.6 -16.4 |

**Figure 5.**
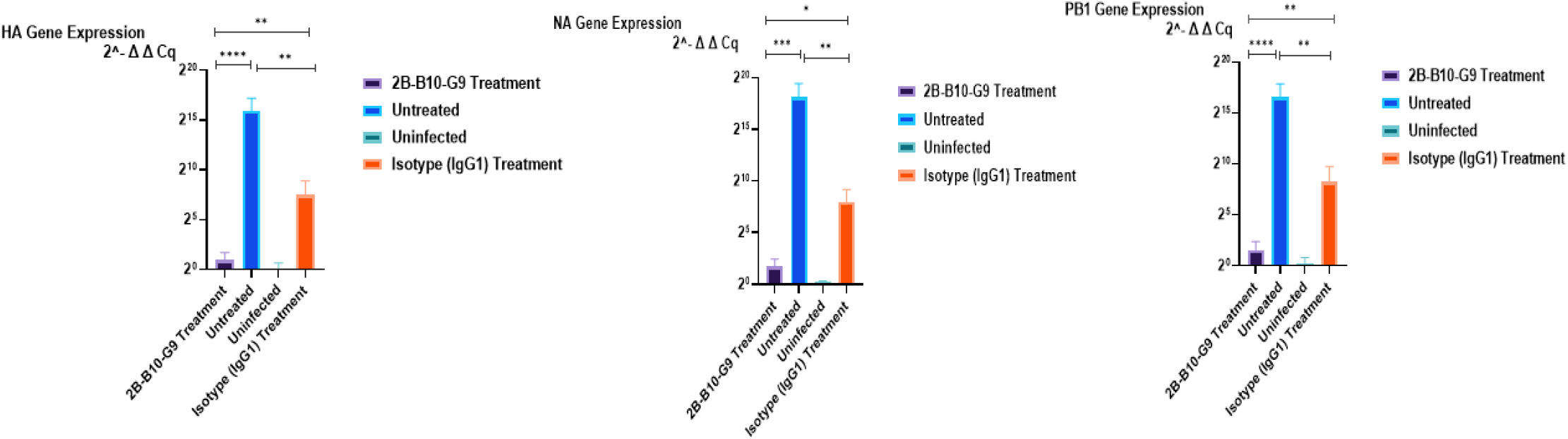
Anti-matrix 1 protein mAb 2B-B10-G9 suppresses virulence gene expressions of H1N1 subtype of influenza A virus. *In ovo* expression of the virulence genes hemagglutinin,(HA) neuraminidase (NA) and polymerase basic protein 1 (PB1) of A/Puerto Rico/8/ 1934 (PR8, H1N1) are depicted in the treatment groups with mAb 2B-B10-G9, untreated, uninfected, and isotype IgG1 mAb treated groups. Experiment was conducted in three biological samples for each group and repeated three times on different days. Data were represented as mean ± SD. One-Way Repeated Measures ANOVA was used to compare the means between the two groups. *p < 0.05, **p < 0.01, ***p < 0.001 and ****P < 0.0001.

## DISCUSSION

Research on novel influenza treatments has primarily focused on broadly reactive mAbs targeting exposed envelope proteins—HA, NA, and M2/BM2 of IAV and IBV.^14–17, 35, 37–39^ While internal proteins like PB1, NP, and the highly abundant M1 are over 90% conserved compared to envelope proteins, mAbs against these internal targets remain largely unexplored. ^22, 40–43^

Previous study showed that anti-M1 mAb 2B-B10-G9 exhibits broad reactivity with the CTD domain of various strains of H1N1, H2N2, and H3N2 subtypes via ELISA.^22^ Our data confirmed heterosubtypic binding of mAb 2B-B10-G9 at the 30 kDa region, with net PI/net AFI ratios for IAV H1N1 and H3N2 subtypes twice as high as for IAV H1N1pdm09 subtype and three to four times higher than for IBV prototype B/Lee/40 and IBV Victoria lineage.

The PRNT is considered the gold-standard methodology and is widely used to evaluate the inhibitory effects of antibodies to viral infections.^29^ mAb 2B-B10-G9 demonstrated 90% reduction in the number of plaques and significant reduction of PFU/ml for both IAV H1N1 and H3N2 subtypes. (p< 0.0001, Figure 4) This result signifies the conserved nature of the M1 protein across the IAV subtypes and importance of the CTD domain in influenza virus life cycle.

Primary cytotoxic mechanisms of mAbs are antibody-dependent cellular cytotoxicity (ADCC) and antibody-dependent cellular phagocytosis (ADCP).^44^ Complement activation by mAbs is considered to contribute to ADCP.^45^ mAbs targeting internal proteins may offer broad immune-mediated protection via cytotoxic T-cell activation. ^46–49^ Studies in embryonated chicken eggs highlight similarities between the avian and human immune systems, particularly in T-cell and dendritic cell functions.^50, 51^ Recent findings indicate that IgG1 mAbs enhance FcγR-dependent effector functions through allosteric changes upon Fab antigen binding.^52^-54 Our study demonstrated the significance of anti-M1 mAb 2B-B10-G9 on the replication of H1N1 subtype of IAV with respect to the isotype IgG1 mAb. (p < 0.01, Figure 5) Fab-mediated immune complex formation and Fc-dependent effector functions were considered to play a significant role in viral clearence with mAb 2B-B10-G9. The isotype IgG1 mAb showed only nonspecific viral clearance most likely via Fc-mediated ADCP and ADCC. The suppression effect of isotype IgG1 mAb on virus replication was less significant with respect to mAb 2B-B10-G9. (Table 4, Figure 5) The 1918 pandemic (H1N1) virus’s extreme virulence was linked to the HA, NA, and PB1 genes.^55^-^57^ We observed a significant *in ovo* reduction in the HA, NA, and PB1 gene expressions of PR8, H1N1 after treatment with 40 µg/ml of 2B-B10-G9, with log2FC of -14.97, -16.44, and -15.02 respectively, compared to the untreated group. A similar trend of reduction in the HA, NA, and PB1 gene expressions was detected with the isotype IgG1 mAb treated group with log2FC of -8.41, -10.15, and -8.29 respectively. However, there was a significant difference of reduction of virulence gene expressions between mAb 2B-B10-G9 and isotype IgG1 mAb treated groups. (p< 0.01) (Figure 5)

## Conclusion

Anti-M1 mAb 2B-B10-G9 exhibits heterosubtypic binding to the M1 protein of IAV H1N1, H1N1pdm09 and H3N2 subtypes, shows *in vitro* neutralization across IAV subtypes, and suppresses the HA, NA and PB1 gene expressions of PR8, H1N1 *in ovo*. This study demonstrated the conserved nature of the M1 protein across the subtypes of IAV, and underlined the significant role of CTD domain of the M1 protein in influenza virus life cycle that makes it a potential therapeutic option for influenza infections. *In vivo* studies in a ferret model will be the subject of future investigations.

## Supporting information

Supplemental MTT Assay

## Acknowledgement

Special thanks to Dr. Ravi Sachidanandam for comments and suggestions.

## REFERENCES

1. Liang Y. Pathogenicity and virulence of influenza. Virulence. 2023 Dec;14(1):2223057. doi: 10.1080/21505594.2023.2223057. PMID: 37339323; PMCID: PMC10283447.

2. Centers for Disease Control and Prevention. Types of Influenza Viruses 2024. https://www.cdc.gov/flu/about/viruses-types Accessed September 18, 2024

3. Centers for Disease Control and Prevention. 2024. Influenza Type A Viruses..

4. Francis, T. J. 1940. A new type of virus from epidemic influenza. Science 92: 405–408.

5. Centers for Disease Control and Prevention. 2024. Trivalent Influenza Vaccines..

6. Bouvier, N. M., and P. Palese. 2008. The biology of influenza viruses. Vaccine 26: D49–D53.

7. Krammer, F., G. J. Smith, R. A. Fouchier, M. Peiris, K. Kedzierska, P. C. Doherty, and others. 2018. Influenza. Nature Reviews Disease Primers 4: 3–24.

8. Wolff, T., and M. Veit. 2021. Influenza B, C and D Viruses (Orthomyxoviridae). In Encyclopedia of Virology (Fourth Edition) D. H. Bamford, and M. Zuckerman, eds. Academic Press, Oxford. 561–574.

9. Saletti, D., Radzimanowski, J., Effantin, G. et al. The Matrix protein M1 from influenza C virus induces tubular membrane invaginations in an in vitro cell membrane model. Sci Rep 7, 40801 (2017).

10. Noton, S. L., E. Medcalf, D. Fisher, A. E. Mullin, D. Elton, and P. Digard. 2007. Identification of the domains of the influenza A virus M1 matrix protein required for NP binding, oligomerization and incorporation into virions. Journal of General Virology 88: 2280–2290.

11. Lampejo, T. 2020. Influenza and antiviral resistance: an overview. European Journal of Clinical Microbiology & Infectious Diseases 39: 1201–1208.

12. Centers for Disease Control and Prevention. 2022. Influenza antiviral drug resistance..

13. Ison, M. G., F. G. Hayden, A. J. Hay, L. V. Gubareva, E. A. Govorkova, E. Takashita, and others. 2021. Influenza polymerase inhibitor resistance: assessment of the current state of the art – a report of the antiviral group. Antiviral Research 194: 105158.

14. Ekiert, D. C., and I. A. Wilson. 2012. Broadly neutralizing antibodies against influenza virus and prospects for universal therapies. Current Opinion in Virology 2: 134–141.

15. Raymond, D. D., G. Bajic, J. Ferdman, P. Suphaphiphat, E. C. Settembre, M. A. Moody, A. G. Schmidt, and S. C. Harrison. 2018. Conserved epitope on influenza-virus hemagglutinin head defined by a vaccine-induced antibody. Proceedings of the National Academy of Sciences 115: 168–173.

16. Yasuhara, A., S. Yamayoshi, M. Kiso, Y. Sakai-Tagawa, M. Okuda, and Y. Kawaoka. 2022. A broadly protective human monoclonal antibody targeting the sialidase activity of influenza A and B virus neuraminidases. Nat Commun 13: 6602.

17. Ramos, E. L., J. L. Mitcham, T. D. Koller, A. Bonavia, D. W. Usner, G. Balaratnam, P. Fredlund, and K. M. Swiderek. 2015. Efficacy and safety of treatment with an anti-M2e monoclonal antibody in experimental human influenza. Journal of Infectious Diseases 211: 1038–1044.

18. Brauer, R., and P. Chen. 2015. Influenza virus propagation in embryonated chicken eggs. J Vis Exp 52421.

19. Reimer, C. B., R. S. Baker, T. E. Newlin, and M. L. Havens. 1966. Influenza virus purification with the zonal ultracentrifuge. Science 152: 1379–1381.

20. Huprikar, J., and S. Rabinowitz. 1980. A simplified plaque assay for influenza viruses in Madin-Darby kidney (MDCK) cells. J Virol Methods 1: 117–120.

21. Reed, L. J., and H. Muench. 1938. A SIMPLE METHOD OF ESTIMATING FIFTY PER CENT ENDPOINTS12. American Journal of Epidemiology 27: 493–497.

22. Bucher, D., S. Popple, M. Baer, A. Mikhail, Y.-F. Gong, E. Whitaker, and others. 1989. M protein (M1) of influenza virus: antigenic analysis and intracellular localization with monoclonal antibodies. Journal of Virology 63: 3622–3633.

23. Köhler, G., and C. Milstein. 1975. Continuous cultures of fused cells secreting antibody of predefined specificity. Nature 256: 495–497.

24. Fishman, J. B., and E. A. Berg. 2019. Protein A and Protein G Purification of Antibodies. Cold Spring Harb Protoc 2019.

25. Lowry, O. H., N. J. Rosebrough, A. L. Farr, and R. J. Randall. 1951. Protein measurement with the Folin phenol reagent. J Biol Chem 193: 265–275.

26. Borejdo, J., and C. Flynn. 1984. Electrophoresis in the presence of Coomassie brilliant blue R-250 stains polyacrylamide gels during protein fractionation. Anal Biochem 140: 84–86.

27. Western Blot Protocols and Recipes - US..

28. 2022. Quantifications of Western Blots with ImageJ / quantifications-of-western-blots-with-imagej.pdf / PDF4PRO. PDF4PRO.

29. https://broadpharm.com/protocol_files/cell_viability_assays. Guidelines for plaque-reduction neutralization testing of human antibodies to dengue viruses..

30. AllPrep DNA/RNA Kits | DNA/RNA Purification Kits | QIAGEN..

31. NanoDrop Eight Spectrophotometer - US..

32. Fulvini, A. A., M. Ramanunninair, J. Le, B. A. Pokorny, J. M. Arroyo, J. Silverman, R. Devis, and D. Bucher. 2011. Gene constellation of influenza A virus reassortants with high growth phenotype prepared as seed candidates for vaccine production. PLoS One 6: e20823.

33. Qin, N., X. Shan, X. Sun, S. Liswaniso, I. Chimbaka, and R. Xu. 2020. Evaluation and Validation of the Six Housekeeping Genes for Normalizing Mrna Expression in the Ovarian Follicles and Several Tissues in Chicken. Braz. J. Poult. Sci. 22: eRBCA-2019-1256.

34. Ruiz-Villalba, A., J. M. Ruijter, and M. J. B. van den Hoff. 2021. Use and Misuse of Cq in qPCR Data Analysis and Reporting. Life (Basel) 11: 496.

35. Ho, K. H., and A. Patrizi. 2021. Assessment of common housekeeping genes as reference for gene expression studies using RT-qPCR in mouse choroid plexus. Sci Rep 11: 3278.

36. Differential gene expression..

37. Guthmiller, J. J., J. Han, H. A. Utset, L. Li, L. Y. Lan, C. Henry, C. T. Stamper, M. McMahon, G. O’Dell, M. L. Fernández-Quintero, A. W. Freyn, F. Amanat, and others. 2022. Broadly neutralizing antibodies target a haemagglutinin anchor epitope. Nature 602: 314–320.

38. Nachbagauer, R., D. Shore, H. Yang, S. K. Johnson, J. D. Gabbard, S. M. Tompkins, J. Wrammert, P. C. Wilson, J. Stevens, R. Ahmed, F. Krammer, and A. H. Ellebedy. 2018. Broadly Reactive Human Monoclonal Antibodies Elicited following Pandemic H1N1 Influenza Virus Exposure Protect Mice against Highly Pathogenic H5N1 Challenge. J Virol 92: e00949–18.

39. Marjuki, H., V. P. Mishin, N. Chai, M. W. Tan, E. M. Newton, J. Tegeris, K. Erlandson, M. Willis, J. Jones, T. Davis, J. Stevens, and L. V. Gubareva. 2016. Human monoclonal antibody 81.39a effectively neutralizes emerging influenza A viruses of group 1 and 2 hemagglutinins. Journal of Virology 90: 10446–10458.

40. Rijnink, W. F., D. Stadlbauer, E. Puente-Massaguer, N. M. Okba, E. Kirkpatrick, S. Strohmeier, and others. 2023. Characterization of non-neutralizing human monoclonal antibodies that target the M1 and NP of influenza A viruses. Journal of Virology 97: e01646–22.

41. Vogel, O. A., and B. Manicassamy. 2020. Broadly Protective Strategies Against Influenza Viruses: Universal Vaccines and Therapeutics. Front Microbiol 11: 135.

42. Van den Hoecke, S., M. Ballegeer, B. Vrancken, L. Deng, E. R. Job, K. Roose, B. Schepens, L. Van Hoecke, P. Lemey, and X. Saelens. 2021. In Vivo Therapy with M2e-Specific IgG Selects for an Influenza A Virus Mutant with Delayed Matrix Protein 2 Expression. mBio 12: e0074521.

43. Carragher, D. M., D. A. Kaminski, A. Moquin, L. Hartson, and T. D. Randall. 2008. A novel role for non-neutralizing antibodies against nucleoprotein in facilitating resistance to influenza virus. J Immunol 181: 4168–4176.

44. Yen, H.-L., E. Hoffmann, G. Taylor, C. Scholtissek, A. S. Monto, R. G. Webster, and E. A. Govorkova. 2006. Importance of neuraminidase active-site residues to the neuraminidase inhibitor resistance of influenza viruses. J Virol 80: 8787–8795.

45. Scholtissek, C., J. Stech, S. Krauss, and R. G. Webster. 2002. Cooperation between the Hemagglutinin of Avian Viruses and the Matrix Protein of Human Influenza A Viruses. J Virol 76: 1781–1786.

46. Presta, L. G. 2006. Engineering of therapeutic antibodies to minimize immunogenicity and optimize function. Adv Drug Deliv Rev 58: 640–656.

47. Hansel, T. T., H. Kropshofer, T. Singer, J. A. Mitchell, and A. J. T. George. 2010. The safety and side effects of monoclonal antibodies. Nat Rev Drug Discov 9: 325–338.

48. Von Holle, T. A., and M. A. Moody. 2019. Influenza and Antibody-Dependent Cellular Cytotoxicity. Front Immunol 10: 1457.

49. Terajima, M., J. Cruz, M. D. T. Co, J.-H. Lee, K. Kaur, J. Wrammert, P. C. Wilson, and F. A. Ennis. 2011. Complement-dependent lysis of influenza a virus-infected cells by broadly cross-reactive human monoclonal antibodies. J Virol 85: 13463–13467.

50. Terajima, M., J. Cruz, A. M. Leporati, L. Orphin, J. A. B. Babon, M. D. T. Co, P. Pazoles, J. Jameson, and F. A. Ennis. 2008. Influenza A virus matrix protein 1-specific human CD8+ T-cell response induced in trivalent inactivated vaccine recipients. J Virol 82: 9283–9287.

51. Padilla-Quirarte, H. O., D. V. Lopez-Guerrero, L. Gutierrez-Xicotencatl, and F. Esquivel-Guadarrama. 2019. Protective Antibodies Against Influenza Proteins. Front Immunol 10: 1677.

52. Kolpe, A., M. Arista-Romero, B. Schepens, S. Pujals, X. Saelens, and L. Albertazzi. 2019. Super-resolution microscopy reveals significant impact of M2e-specific monoclonal antibodies on influenza A virus filament formation at the host cell surface. Sci Rep 9: 4450.

53. Garcia, P., Y. Wang, J. Viallet, and Z. Macek Jilkova. 2021. The Chicken Embryo Model: A Novel and Relevant Model for Immune-Based Studies. Front Immunol 12: 791081.

54. Orlandi, C., D. Deredge, K. Ray, N. Gohain, W. Tolbert, A. L. DeVico, P. Wintrode, M. Pazgier, and G. K. Lewis. 2020. Antigen-Induced Allosteric Changes in a Human IgG1 Fc Increase Low-Affinity Fcγ Receptor Binding. Structure 28: 516-527.e5.

55. Tumpey, T. M., A. García-Sastre, J. K. Taubenberger, P. Palese, D. E. Swayne, and C. F. Basler. 2004. Pathogenicity and immunogenicity of influenza viruses with genes from the 1918 pandemic virus. Proc Natl Acad Sci U S A 101: 3166–3171.

56. Lamb, R. A., C.-J. Lai, and P. W. Choppin. 1981. Sequences of mRNAs derived from genome RNA segment 7 of influenza virus: \ colinear and interrupted mRNAs code for overlapping proteins. Proceedings of the National Academy of Sciences 78: 4170–4174.

57. Dubois, J., O. Terrier, and M. Rosa-Calatrava. 2014. Influenza Viruses and mRNA Splicing: Doing More with Less. mBio 5: e00070–14.

