## Supplemental MTT Assay for "Anti-Matrix Protein 1 Monoclonal Antibody Neutralizes Influenza A Virus Subtypes": AMP1Supplement.docx.html

 

MTT Assay for Anti-matrix protein 1 mAb 2B-B10-G9 shows non-significant reduction of MDCK cell viability at 40 ug/ml.

MTT Assay for mAb 2B-B10-G9 showed non-significant reduction of the MDCK cell viability between 0.5 ug/ml and 40 ug/ml with respect to DMSO vehicle control (Figure 1). The reduction of MDCK cell viability was significant at doses of 100 ug/ml and 200 ug/ml. (p < 0.0001) (Figure 1). The therapeutic dose of mAb 2B-B10-G9 for in vitro and in ovo studies was determined as 40 ug/ml according to the MTT assay.

Figure 1. MTT Assay for mAb 2B-B10-G9 at doses of 0.5, 1.0, 5.0, 10.0, 40.0, 100.0 and 200.0 ug/ml with negative (vehicle) control Dimethyl sulfoxide (DMSO) and positive control apoptotic agent Staurosporin. Data (n =3) are represented as mean ± S.D. One-Way ANOVA Multiple Comparisons was used to compare the means.  ns: non-significant,  \*\*\*\*p< 0.0001
