## Supplementary figures and images for "Anti-Matrix Protein 1 Monoclonal Antibody Neutralizes Influenza A Virus Subtypes"

### image1.png

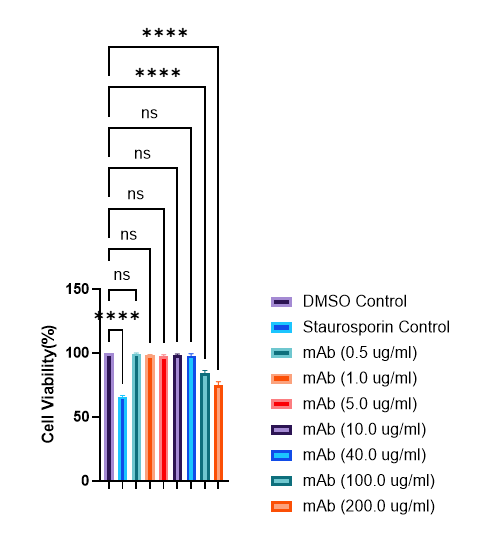
